# The Atypical Rho GTPase CHW-1 Works With SAX-3/Robo to Mediate Axon Guidance in *Caenorhabditis elegans*

**DOI:** 10.1101/265066

**Authors:** Jamie K. Alan, Sara Robinson, Katie Magsig, Rafael S. Demarco, Erik A. Lundquist

## Abstract

During development, neuronal cells extend an axon towards their target destination in response to a cue to form a properly functioning nervous system. Rho proteins, Ras-related small GTPases that regulate cytoskeletal organization and dynamics, cell adhesion, and motility, are known to regulate axon guidance. Despite extensive knowledge about canonical Rho proteins (RhoA/Rac1/Cdc42), little is known about the *Caenorhabditis elegans* (*C. elegans*) atypical Cdc42-like family members CHW-1 and CRP-1 in regards to axon pathfinding and neuronal migration. *chw-1*(Chp/Wrch) encodes a protein that resembles human Chp (Wrch-2/RhoV) and Wrch-1 (RhoU), and *crp-1* encodes for a protein that resembles TC10 and TCL. Here, we show that *chw-1* works redundantly with *crp-1* and *cdc-42* in axon guidance. Furthermore, proper levels of *chw-1* expression and activity are required for proper axon guidance. When examining CHW-1 GTPase mutants, we found that the native CHW-1 protein is likely partially activated, and mutations at a conserved residue (position 12 using Ras numbering, position 18 in CHW-1) alter axon guidance and neural migration. Additionally, we showed that *chw-1* genetically interacts with the guidance receptor *sax-3* in PDE neurons. Finally, in VD/DD motor neurons, *chw-1* works downstream of *sax-3* to control axon guidance. In summary, this is the first study implicating the atypical Rho GTPases *chw-1* and *crp-1* in axon guidance. Furthermore, this is the first evidence of genetic interaction between *chw-1* and the guidance receptor *sax-3*. These data suggest that *chw-1* is likely acting downstream and/or in parallel to *sax-3* in axon guidance.

## INTRODUCTION

During proper development of the nervous system, neurons must first migrate to their final position where they then extend axons in order to establish synaptic connections. In order to first migrate and then extend an axon, the neuronal cells must undergo dynamic changes in their actin cytoskeleton migration (Tessier-Lavigne and Goodman 1996) (Dickson 2002; Gallo and Letourneau 2002). Many studies have shown that Rho proteins are required for actin cytoskeleton rearrangement and subsequent neuronal migration and axon extension (Lundquist 2003; Luo 2000; Dickson 2001). Rho family proteins are Ras-related small GTPases that regulate cytoskeletal organization and dynamics, cell adhesion, motility, trafficking, proliferation, and survival (Etienne-Manneville and Hall 2002). Misregulation of Rho proteins can result in defects in cell proliferation, cell adhesion, cell morphology, and cell migration (Ridley 2004). These proteins function as tightly regulated molecular switches, cycling between an active GTP-bound state and an inactive GDP-bound state (Jaffe and Hall 2005). Despite extensive knowledge about regulation and function of the canonical Rho proteins (RhoA/Rac1/Cdc42), little is known about the contribution of the *Caenorhabditis elegans (C. elegans*) atypical Cdc42-like family members CHW-1 and CRP-1 to axon pathfinding and neuronal migration. *C. elegans* contains a single gene, *chw-1* which encodes a protein that resembles human Chp (Wrch-2/RhoV) and Wrch-1 (RhoU), and *crp-1* encodes for a protein that resembles TC10 and TCL.While *chw-1* has not been studied in the context of axon pathfinding, it has been implicated in cell polarity, working with LIN-18/Ryk and LIN-17/Frizzled(Kidd et al. 2015). Additionally, there is a body of literature concerning the related mammalian homologues Wrch-1 (Wnt-regulated Cdc42 homolog-1, RhoU) and Chp (Wrch-2/RhoV), the most recently identified Rho family members. Wrch-1 and Chp share 57% and 52% sequence identity with Cdc42, respectively, and 61% sequence identity with each other (Tao et al. 2001; Aronheim et al. 1998). Wrch-1 and Chp are considered atypical GTPases for a number of reasons. Atypical Rho proteins vary in either their regulation of GTP/GDP-binding, the presence of other domains besides the canonical Rho insert domain, and variances in their N- and C-termini and/or posttranslational modifications. When compared to the other members of the Cdc42 family, Wrch-1 has elongated N-terminal and C-terminal extensions. The N-terminus of Wrch-1 has been shown to be an auto-inhibitory domain, and removal of the N-terminus increases the biological activity of the protein (Shutes et al. 2004; Shutes et al. 2006; Berzat et al. 2005). The N-termini of both Wrch-1 and Chp contain PxxP motifs, which may mediate interactions with proteins containing SH3 domains, such as the adaptor proteins Grb2 and Nck (Zhang et al.). Wrch-1 is also divergent in the C-terminus, when compared to other members of the Cdc42 family. Like Chp (Chenette et al. 2006), instead of being irreversibly prenylated on a CAAX motif, Wrch-1 is reversibly palmitoylated on a CFV motif (Berzat et al. 2005). This reversible modification may lead to more dynamic regulation of the protein.

Although Wrch-1 shares sequence identity with Cdc42 and Chp-1, Wrch-1 shares only partially overlapping localization and effector interactions with these proteins, and its activity is regulated in a distinct manner. Like Cdc42, Wrch-1 activity leads to activation of PAK1 and JNK (Chuang et al. 2007; Tao et al. 2001), formation of filopodia (Ruusala and Aspenstrom 2008; Saras et al. 2004), and both morphological (Brady et al. 2009) and growth transformation in multiple cell types (Berzat et al. 2005; Brady et al. 2009). In addition, Wrch-1 also regulates focal adhesion turnover (Chuang et al. 2007; Ory et al. 2007), negatively regulates the kinetics of tight junction formation (Brady et al. 2009), plays a required role in epithelial morphogenesis (Brady et al. 2009), modulates osteoclastogenesis (Brazier et al. 2009; Ory et al. 2007; Brazier et al. 2006), and regulates neural crest cell migration (Faure and Fort 2011). Chp/Wrch-2, a protein highly related to Wrch-1, was identified in a screen that was designed to look for proteins that interact with p21-activated kinase (Pak1) (Aronheim et al. 1998). Like Wrch-1, Chp also contains N- and C-terminal extensions when compared to Cdc42 and the N-terminus auto-inhibitory domain (Shutes et al. 2004; Chenette et al. 2005). Also like Wrch-1, active Chp leads to cell transformation (Chenette et al. 2005), signals through PAK (PAK6) (Shepelev and Korobko 2012) and JNK (Shepelev et al. 2011), and is involved in neural crest cell specification and migration (Faure and Fort 2011; Guemar et al. 2007).

While the GTPase domain of CHW-1 is highly similar to human Wrch-1 and Chp/Wrch-2, it is divergent in its N- and C-terminus. The N-terminal extension found in both Wrch-1 and Chp/Wrch-2, which regulates the function of these proteins, is absent in CHW-1. Additionally, CHW-1 lacks any C-terminal lipid modification motif, such as a CXX motif found in Wrch-1 and Chp/Wrch-1, suggesting that it does not require membrane targeting for its function. While CHW-1 bears some sequence identity to both Wrch-1, Chp/Wrch-1 and Cdc42, it is most similar to human Wrch-1(Kidd et al. 2015).

The *C. elegans* protein CRP-1, a member of the Cdc42 subfamily, shares sequence identity, possess atypical enzymatic activity and possesses similar functions to the mammalian GTPases TC10 and TCL (Aspenstrom et al. 2004; de Toledo et al. 2003; Michaelson et al. 2001; Murphy et al. 2001; Murphy et al. 1999; Jenna et al. 2005; Vignal et al. 2000). Interestingly, like CRP-1, TC10 and TCL localize at internal membranes (Michaelson et al. 2001; Aspenstrom et al. 2004; Vignal et al. 2000; Jenna et al. 2005). Also, like CRP-1, TC10 and TCL mediate cellular trafficking (Jenna et al. 2005; Okada et al. 2008; de Toledo et al. 2003). However, unlike the rest of the Cdc42 family members, overexpression of CRP-1 was unable to induce cytoskeletal changes in mouse fibroblasts, suggesting that CRP-1 may have overlapping and unique functions compared to other Cdc42 family members (Jenna et al. 2005). Indeed, unlike other Cdc42 family members, *crp-1* is an essential component for UPR-induced transcriptional activities. *crp-1* mediates this processes through physical and genetic interactions with the AAA+ ATPase CDC-48 (Caruso et al. 2008).

Because members of the Rho family have many overlapping functions, in addition to their unique functions, we hypothesized that atypical Rho GTPases, like the other Rho family small GTPases, may contribute to axon guidance. To test this hypothesis, we genetically characterized the *C. elegans* atypical Rho GTPases *chw-1* and *crp-1*, in axon pathfinding. We found the *chw-1* works redundantly with *crp-1* and *cdc-42* in axon guidance. Next, we examined CHW-1 GTPase mutants. Most GTPases have a conserved glycine at position 12 (using Ras numbering). However, CHW-1 possesses an alanine at this conserved position (position 18 in CHW-1), which is predicted to be partially activating. We found that ectopic expression of either CHW-1, CHW-1(A18V) and CHW-1(A18G) caused axon guidance defects, suggesting proper expression and activity levels of CHW-1 are required for proper axon guidance. The A18V mutation is predicted to be activating, and the A18G mutation is predicted to reduce function. Furthermore, we observed that an activated version of CHW-1 (CHW-1(A18V)) resulted in formation of ectopic lamellipodia, similar to activated versions of other Rho proteins such as MIG-2, CED-10, and CDC-42. These data suggest that, like other Rho GTPases and their mammalian counterparts, CHW-1 is likely modulating the actin cytoskeleton to regulate axon pathfinding. When examining the CHW-1(A18G) mutant, we found that it displayed anteriorly displaced amphid neuron cell bodies. This phenotype is similar to a loss-of-function *sax-3* phenotype. Therefore, we hypothesized that *chw-1* may genetically interact with *sax-3* in axon guidance. Indeed, we found that *chw-1* does genetically interact with *sax-3* in the amphid and PDE neurons. Furthermore, we were able to show that loss of *chw-1* attenuates axon pathfinding defects driven by activated SAX-3 in VD/DD motor neurons. These data suggest that *chw-1* is likely acting downstream and/or in parallel to *sax-3* in axon guidance.

In summary, this is the first study characterizing the role of the *C. elegans* atypical Rho GTPases in axon guidance. We found that *chw-1* is acting redundantly with *crp-1* and *cdc-42* in axon guidance. Like other Rho GTPases, CHW-1 is involved in axon guidance, and it works redundantly with CRP-1 and CDC-42 in this context. Furthermore, we showed that *chw-1* interacts genetically with the guidance receptor *sax-3* in axon guidance.

## MATERIALS AND METHODS

### C. elegans genetics and transgenics

*C. elegans* were cultured using standard techniques (Brenner 1974; J and J 1988). All of the experiments were done at 20°C, unless otherwise noted. Transgenic worms were made by gonadal micro-injection using standard techniques (Mello and Fire 1995). RNAi was done using clones from the Ahringer library, as previously described (Kamath and Ahringer 2003). The following mutations and constructs were used in this work: LGI- *unc-40(n324);* LGII- *cdc-42(gk388), mIn1;* LGV- *chw-1(ok697), crp-1(ok685);* LGX- *sax-3(ky123), unc-6(ev400), IqIs2 (osm-6::gfp);* LG unassigned- *osm-6::chw-1::gfp, osm-6::chw-1(A18G)::gfp, osm-6::chw-1(A18V)::gfp, unc- 25::myr::sax-3.*

The mutants were maintained as homozygous stocks, when possible. In some cases, this was not possible because the double mutants were lethal or maternal-lethal, and thus could not be maintained as homozygous stocks. If this was the case, the mutation was maintained in a heterozygous state over a balancer chromosome. *cdc-42(gk388*) was balanced by *mIn1, which* harbors a transgene that drives *gfp* expression in the pharynx. *cdc-42(gk388*) homozygotes were identified by lack of pharyngeal green fluorescent protein (GFP).

### Scoring of PDE and VD/DD axon defects

Axon pathfinding defects were scored in the fourth larval stage (L4) or in young pre-gravid adult animals harboring green fluorescent protein (gfp), which was expressed in specific cell types. In the event that the animals did not survive to this stage, they were counted at an earlier stage with appropriate controls. The PDE neurons and axons were visualized in cells that had an *osm-6::gfp* transgene (*lqIs2 X*), which is expressed in all ciliated neurons including the PDE. A PDE axon was considered to be misguided if it failed to extend an axon directly to the ventral nerve cord (VNC). If the trajectory of the axon was greater than a 45° angle from the wild-type position, the axon was scored as misguided, whether it eventually reached the VNC or not. The neurons were considered to have ectopic lamellipodia if they had a lamellipodia-like structure protruding from anywhere on the cell body, axon, or dendrite.

The VD/DD motor neuron morphology was scored in animals that had an *unc- 25::gfp* transgene (*juIs76* II)(Jin et al. 1999). The *unc-25* promoter is turned on in all GABAergic neurons, including the VDs and DDs. In wild-type animals, the VD/DD commissures extend directly from the VNC toward the dorsal surface of the animals. At the dorsal surface, they form the dorsal nerve cord. Aberrant VD/DD pathfinding was noted when the commissural axons were misguided or terminated prematurely.

### Molecular biology

The CHW-1 constructs were subcloned from cDNA in a pBS (bluescript) vector backbone, which were a generous gift from Dr. Ambrose Kidd, who made these constructs while in the laboratories of Dr. Adrienne Cox and Dr. Channing Der (UNC-Chapel Hill) in collaboration with Dr. Dave Reiner (Texas A&M). Recombinant DNA, polymerase chain reaction (PCR), and other molecular biology techniques were preformed using standard procedures (Sambrook J. 1989). All primer sequences used in the PCR reactions are available upon request.

### Microscopy and imaging

The animals used for the imaging analyses were mounted for microscopy in a drop of M9 buffer on an agarose pad (J and J 1988). Both the buffer and the agarose pad contained 5 mM sodium azide, which was used as an anesthetic. Then, a coverslip was placed over the sample, and the slides were analyzed by epifluorescence for GFP. A Leica DMRE microscope with a Qimaging Rolera MGi EMCCD camera was used, along with Metamorph and ImageJ software.

### Data availability

Strains are available upon request.

## RESULTS

### *chw-1* acts redundantly with *cdc-42* and *crp-1* in axon guidance

As previously described (Kidd et al. 2015), the *C. elegans* genome contains a single gene, F22E12.1, which encodes CHW-1, the *C. elegans* ortholog of human Chp/Wrch-1. The core effector domain of CHW-1 is most similar to human Wrch-1, and is considerably variant from human Chp. The N- and C-termini are considerably different than either human ortholog. While both Wrch-1 and Chp have an N-terminal extension that negatively regulates the protein, this domain is almost entirely absent in Chw-1 (Figure 1). While most Rho proteins terminate in a CAAX motif that signals for an irreversible prenyl lipid modification, both Wrch-1 and Chp end in a CXX motif and are modified by a reversible palmitate modification, which is required for their membrane localization and biological activity (Berzat et al. 2005; Chenette et al. 2005). However, CHW-1 lacks a CAAX or a CXX motif and presumably does not require membrane targeting for its function (Figure 1). Given these differences, we sought to determine the role of *chw-1* in axon guidance. The human orthologs of CHW-1 are in the Cdc42 family of Rho proteins, and in mammalian systems, the Cdc42 family of proteins (Cdc42, Wrch-1, Chp, TC10, and TCL) work in concert to carry out both unique and redundant functions. While CHW-1 is the *C. elegans* ortholog of Wrch-1 and Chp, CRP-1 closely resembles TC10 and TCL in both sequence and function (Caruso et al. 2008; Jenna et al. 2005). Therefore, we first sought to determine whether *chw-1* works redundantly with *crp-1* and/or *cdc42* in axon guidance. For these assays, we visualized the morphology and position of the PDE axons. *C. elegans* has two bilateral PDE neurons that are located in the post-deirid ganglia of the animal. In the wild-type N2 strain, PDE neurons extend their axons ventrally in a straight line from the cell body to the ventral nerve cord. When the axon reaches the ventral cord, it then branches and extends towards both the anterior and the posterior. To visualize the neurons, we used a transgene gene (*lqIs2; osm-6::gfp*) expressed in the PDE neurons. The *osm-6* promoter is expressed in all ciliated sensory neurons including the PDE neurons (Collet et al. 1998). We observed that alone, deletion alleles or RNAi against *chw-1, crp-1*, or *cdc-42* produced very few axon guidance defects (Figure 2). However, when two or more of these genes were knocked out by either a deletion allele or RNAi we observed a synergistic increase in axon guidance defects (up to 50%). Furthermore, when all 3 genes were knocked out the axon guidance defects increased to 70%. Taken together, these genetic data suggest that *chw-1, crp-1*, and *cdc42* act redundantly in PDE axon guidance.

**Figure 1.**
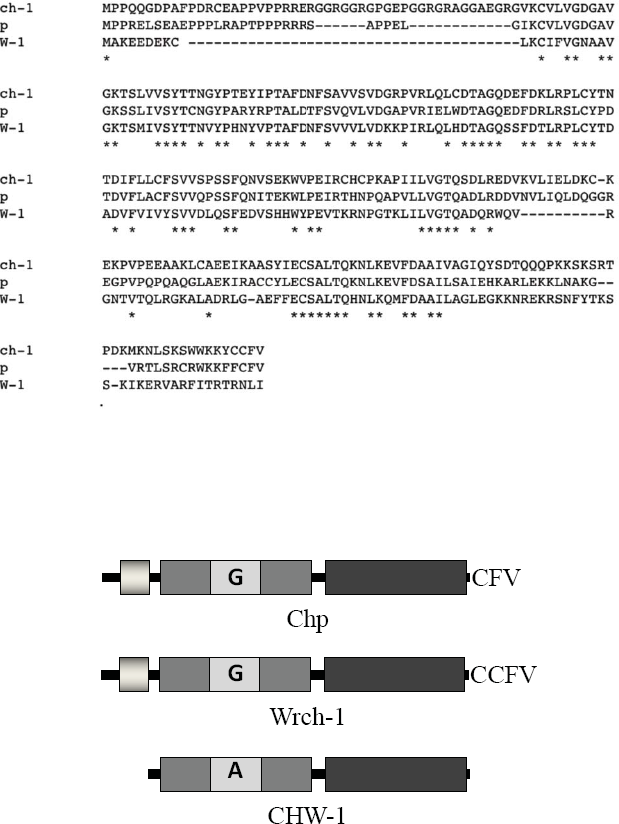
CHW-1 is similar to Wrch-1/RhoU and Chp/Wrch-2/RhoV. Sequence alignment of CHW-1, human Wrch-1, and human Chp done in biology workbench. Identical residues are marked with an asterisk (*). B) CHW-1 lacks the N-terminal extension found in Wrch-1 and Chp (light gray box). CHW-1 also contains an atypical residue (alanine instead of glycine) at position 18 (analogous to position 12 in Cdc42 and most Rho and Ras family members).

**Figure 2.**
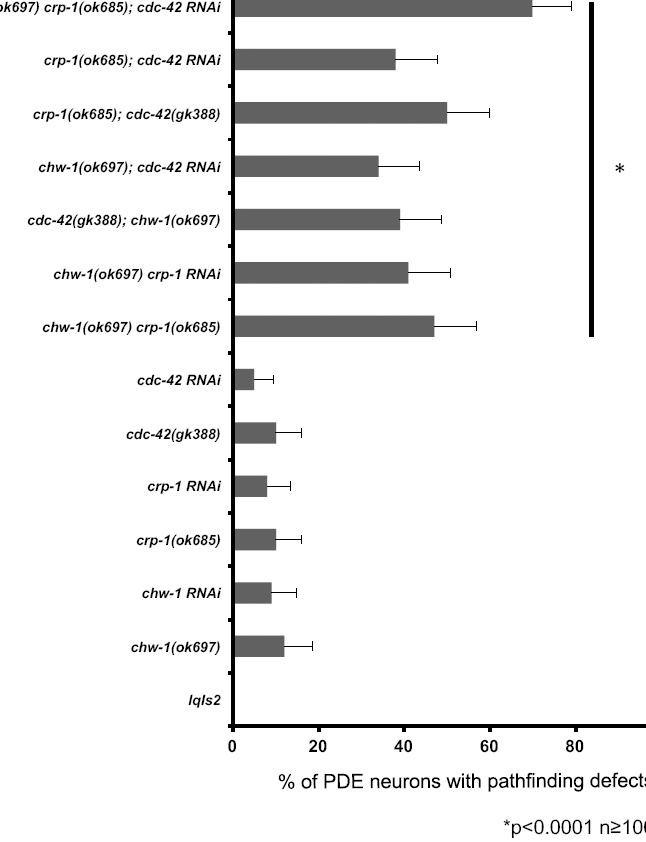
*chw-1* works redundantly with *cdc-42* and *crp-1* in axon guidance. Quantitation of PDE defects. *lqIs2* is the *osm-6::gfp* control transgene. At least 100 neurons were scored for each genotype. p<0.0001 as determined by Fisher Exact Analysis (Graphpad). The error bars represent the standard proportion of the mean.

### Proper expression and activity levels of CHW-1 are required for axon guidance

Ras superfamily GTPases, including the Rho-family GTPases, cycle between active GTP-bound forms and inactive GDP bound forms driven by the intrinsic GTPase activity of the molecules. The glycine 12 to valine (G12V) substitution has been previously shown to result in inhibition of the intrinsic GTPase activity of the GTPase, favoring the active, GTP-bound state (Wittinghofer et al. 1991). The G12V mutation also makes the protein insensitive to GTPase activating proteins (GAPs) (Ahmadian et al. 1997), rendering the protein constitutively GTP-bound and active. Most Rho proteins harbor a glycine at position 12, however CHW-1 possesses an alanine at this conserved position (position 18 in CHW-1), which is predicted to be a partially activating, resulting in greater GTP-bound levels and an increase in activity. Indeed, evidence from other labs suggests that the native CHW-1 protein may be partially constitutively active (Kidd et al. 2015). We found that transgenic expression of CHW-1, CHW-1(A18V), and CHW-1(A18G) all caused axon guidance defects to a very similar degree (24%, 23%, and 29%, respectively) (Figure 3). This evidence suggests that, like the mammalian counterpart Wrch-1, proper levels of CHW-1 activity are required for correct axon guidance (Alan et al.; Brady et al. 2009). In other words, too much or too little activity of CHW-1 results in an aberrant outcome (misguided axons).

**Figure 3.**
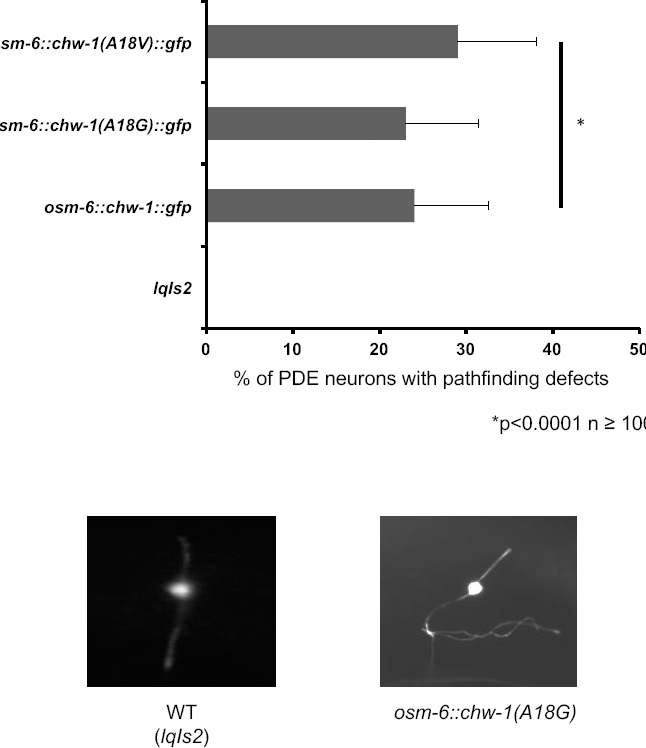
Proper expression and activity levels of CHW-1 are required for axon guidance. A) Quantitation of PDE defects. *lqIs2* is the *osm-6::gfp* control transgene. At least 100 neurons were scored for each genotype. p<0.0001 as determined by Fisher Exact Analysis (Graphpad). The error bars represent the standard proportion of the mean. B) A micrograph of a PDE neuron of a WT adult animal (left panel) and a PDE neuron of an adult animal expressing CHW-1(A18G) (right panel). The scale bar represents 5 μm.

### Expression of CHW-1(A18V) results in the formation of ectopic lamellipodia in PDE neurons

Previously, our lab showed that expression of the Rac GTPases CED-10 and MIG-2 harboring the G12V mutation in neurons *in vivo* resulted in the formation of ectopic lamellipodial and filopodial protrusions (Struckhoff and Lundquist 2003), and that CDC-42(G12V) expression caused similar lamellipodial and filopodial protrusions (Demarco and Lundquist). Therefore, we sought to determine whether a similar mutation at this conserved residue, CHW-1(A18V), also resulted in ectopic lamellipodia. Indeed, we found that expression of CHW-1(A18V) resulted in ectopic lamellipodia in 32% of PDE neurons (Figure 4). Expression of wild type CHW-1 or CHW-1(A18G) did not show ectopic lamellipodia. This suggests that like CDC42 and the Rac GTPases, CHW-1 may be involved in regulation of protrusion during neuronal development. Furthermore, these data suggest that full activation by a G12V mutation is necessary for this lamellipodial protrusion.

**Figure 4.**
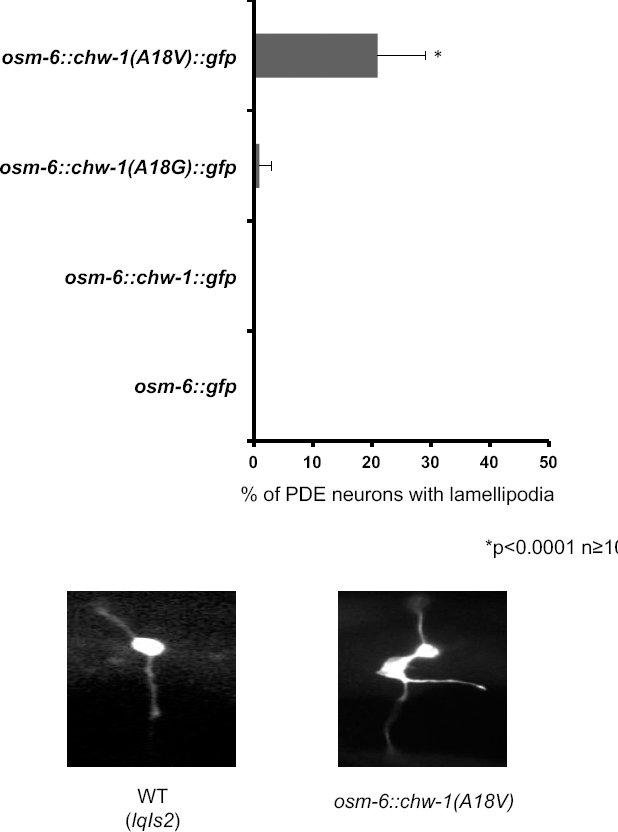
Expression of CHW-1(A18V) results in the formation of ectopic lamellipodia in PDE neurons. A) Quantitation of PDE defects. *osm-6::gfp* is the control transgene. At least 100 neurons were scored for each genotype. p<0.0001 as determined by Fisher Exact Analysis (Graphpad). The error bars represent the standard proportion of the mean. B) A micrograph of a PDE neuron of a WT adult animal (left panel) and a PDE neuron of an adult animal expressing CHW-1(A18V) (right panel).

### Expression of CHW-1(A18G) results in amphid neuron defects

The *osm- 6::gfp* transgene is expressed in the PDE neurons as well as the sensory amphid neurons in the head and the phasmid neurons in the tail. When examining the PDE neurons in the CHW-1(A18G) mutant, we noticed that the morphology of the amphid neurons in the head was altered. The cell bodies of the amphid neurons in the CHW-1(A18G) transgenic animals were displaced anteriorly (Figure 5). Expression of wild-type CHW-1 or CHW-1(A18V) did not cause anterior amphid displacement. Interestingly, although the *osm-6::chw-1(A18G*) transgene displayed defects in amphid neuron morphology, the loss of function *chw-1(ok697*) allele did not display these defects. The *chw-1(ok697) crp-1(ok685*) double mutant also so showed no amphid neuron defects (Figure 5). The endogenous amino acid at this regulatory position in CHW-1 is an alanine, which is predicted to be a partially activating mutation, suggesting that CHW- 1(A18G) may be acting as a dominant-negative in this context.

**Figure 5.**
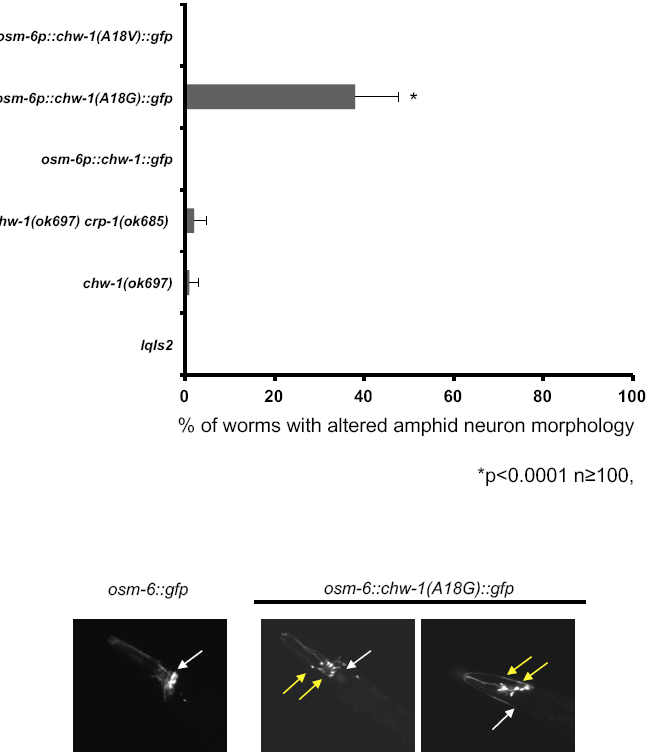
Expression of CHW-1(A18G) results in amphid neuron defects. A) Quantitation of amphid neuron defects. *lqIs2* is the *osm-6::gfp* control transgene.. The Y-axis denotes the genotype and the X-axis represents the percentage of amphid neuron defects. At least 100 neurons were scored for each genotype. p<0.0001 as determined by Fisher Exact Analysis (Graphpad). The error bars represent the standard proportion of the mean. B) A micrograph of the amphid neurons of a WT adult animal (left panel), and the amphid neurons of adult animals expressing CHW-1(A18G) (right two panels). The wild-type position of the amphid neuron cell bodies is indicated by the white arrow and the mutant positions are indicated by the yellow arrows

### *chw-1(ok697*) in a *sax-3(ky123*) background increases amphid neuron defects

We noted that amphid neuron phenotype of CHW-1(A18G) was similar to what was reported for a loss-of-function *sax-3* phenotype (Zallen et al. 1999). *sax-3* encodes an immunoglobulin guidance receptor similar to vertebrate and *Drosophila* Robo. We next investigated whether CHW-1 interacts genetically with SAX-3. First, we examined the amount of amphid neuron defects in the loss-of-function *sax-3(ky123*) animals. We found that 50% of the *sax-3(ky123*) animals had altered amphid neuron morphologies (Figure 6), similar to what was previously reported in the literature (Zallen et al. 1999). The *sax-3(ky123); chw-1(ok697*) double mutants displayed a synergistic increase in the amount of amphid neuron defects (Figure 6), suggesting redundant function. While the *sax-3(ky123*) allele is reported to be a null, these animals are viable (unlike other *sax-3* null alleles), suggesting that the *sax-3(ky123*) allele may retain a small amount of functionality. Thus, we cannot rule out that SAX-3 and CHW-1 might act in the same pathway. We did not observe any genetic interactions between *chw-1* and other axon guidance genes such as *unc-6* and *unc-40.*

**Figure 6.**
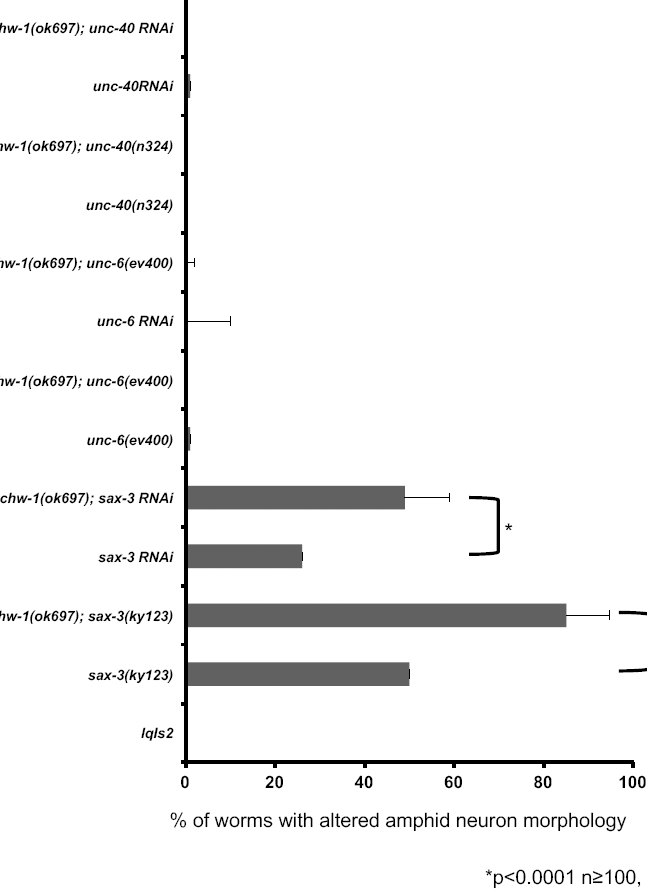
Loss of both *chw-1 and sax-3*, but not other axon guidance molecules, synergistically increases amphid neuron defects. Quantitation of amphid neuron defects. *lqIs2* is the *osm-6::gfp* control transgene. The Y-axis denotes the genotype and the X-axis represents the percentage of amphid neuron defects. At least 100 neurons were scored for each genotype. p<0.0001 as determined by Fisher Exact Analysis (Graphpad). The error bars represent the standard proportion of the mean.

### Loss of both *chw-1* and *sax-3* increases PDE axon pathfinding defects

Next, we sought to determine whether *chw-1* and *sax-3* interacted genetically in the context of PDE axon guidance. *chw-1(ok697*) and *sax-3(ky123*) alone resulted in PDE axon guidance defects (10% and 40%, respectively) (Figure 7). The double mutant (*chw-1(ok697); sax-3(ky123)*) displayed an increase in axon pathfinding defects (68%), suggesting that these genes act redundantly in axon pathfinding. We confirmed this result using RNAi against *sax-3*. Notably, *chw-1* showed no significant interactions with *unc-6* or *unc-40*, genes also involved in ventral PDE axon guidance.

**Figure 7.**
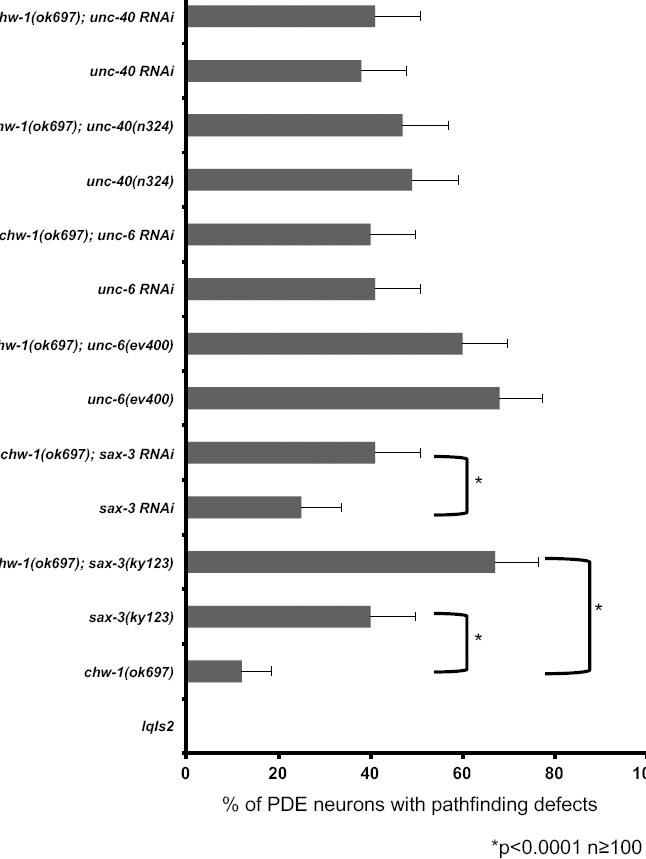
Loss of both *chw-1* and *sax-3* increases PDE axon pathfinding defects. Quantitation of PDE defects. *lqIs2* is the *osm-6::gfp* control transgene. The Y-axis denotes the genotype and the X-axis represents the percentage of ectopic lamellipodia formation. At least 100 neurons were scored for each genotype. *p<0.0001 as determined by Fisher Exact Analysis. The error bars represent the standard proportion of the mean.

### Loss of *chw-1* but not *crp-1* attenuates the effects of *myr::sax-3*

Next, we wanted to further define the relationship between CHW-1 and SAX-3. To do this, we constructed a transgene that expressed the cytoplasmic domain of SAX-3 with an N-terminal myristoylation site. In UNC-40/DCC and UNC-5 receptors, a similar myristoylated construct leads to constitutive activation of these receptors (Gitai et al. 2003; Norris and Lundquist 2011; Norris et al. 2014). The MYR::SAX- 3 transgene did not display a phenotype in the PDE neurons or in the amphid neurons when driven from the *osm-6* promoter. Therefore, we expressed *myr::sax-3* in the VD/DD motor neurons using the *unc-25* promoter. The VD/DD motor neurons are born in the ventral nerve cord and have axons that extend to the dorsal nerve cord. Dorsal guidance of these commissures is not dependent upon endogenous SAX-3 (Li et al. 2013), however activation of SAX-3 in the neurons significantly alters dorsal axon guidance. We observed that MYR::SAX-3 caused axon pathfinding defects in the majority of VD/DD motor neurons (98%) (Figure 8). Loss of *chw-1* significantly attenuated these guidance defects (68%), whereas loss of *crp-1* did not. These results were confirmed using RNAi, where loss of *chw-1* by RNAi also decreased the axon pathfinding defects in the MYR::SAX-3 background (79%) These data suggest that CHW-1 acts with SAX-3 in axon guidance in the VD/DD motor neurons.

**Figure 8.**
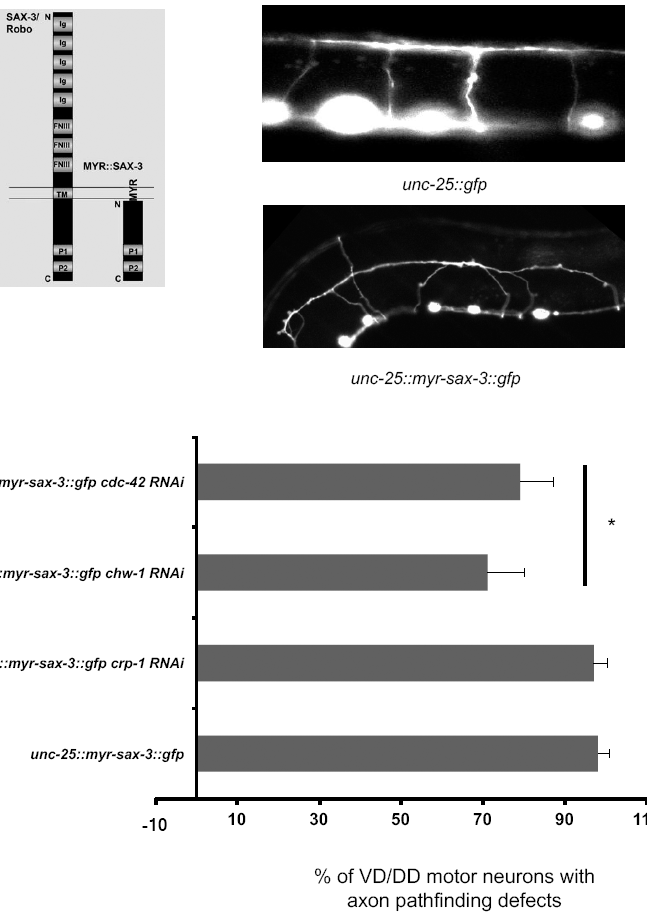
Loss of *chw-1* but not *crp-1* attenuates the defects in the VD/DD motor neurons driven by activated SAX-3. A) A representation of the MYR-SAX-3 construct. B) A micrograph of the VD/DD neurons of a WT adult animal (top panel), and the VD/DD neurons of an adult animal expressing MYR-SAX-3 (bottom panel). C) Quantitation of VD/DD defects. *unc-25::myr-sax-3* is the activated SAX-3 receptor driven by the *unc-25* promoter, which drives expression in the VD/DD motor neurons. The Y-axis denotes the genotype and the X-axis represents the percentage of ectopic lamellipodia formation. The number of axons scored was greater than 100. *p<0.0004 as determined by Fisher Exact Analysis. The error bars represent the standard proportion of the mean.

## DISCUSSION

There have been several studies focused on molecules that function in axon guidance. However, the entire picture of how these molecules function together in concert is still unclear. CDC-42 and other Rho GTPases, such as CED-10 and MIG-2, have been previously characterized in axon guidance (Gallo and Letourneau 1998; Hall and Lalli; Luo 2000). However, there have been no studies describing if and how the other *C. elegans* CDC-42 family members, CHW-1 and CRP-1 contribute to axon guidance. Here, we present evidence suggesting that *chw-1* works redundantly with *crp-1* and *cdc-42* in axon guidance. Furthermore, overexpression of either CHW-1 or CHW-1 GTPase mutants alters axon guidance, suggesting that like the human counterpart Wrch-1, proper levels of CHW-1 expression and activity are required for normal CHW-1 function. For example, in cell culture either knockdown, overexpression or alterations in the GTPase activity of Wrch-1 results in similar disruptions in epithelial cell morphogenesis (Alan et al.; Brady et al. 2009). Taken together, these data suggest that tightly regulated expression and activity levels of CHW-1 and Wrch-1 are required for their normal biological functions.

Next, we showed that a GTPase deficient (activated) mutant of CHW-1, CHW- 1(A18V), results in the formation of ectopic lamellipodia in the PDE neuron. This result is similar to the phenotype observed for activated versions of MIG-2, CED- 10, and CDC-42 (Struckhoff and Lundquist 2003) (Demarco and Lundquist), suggesting that, like these canonical GTPases, CHW-1 regulates the actin cytoskeleton. There is evidence that, similar to CDC-42, Wrch-1 also regulates the actin cytoskeleton and is able to regulate the formation of filopodia in cells (Aspenstrom et al. 2004; Ruusala and Aspenstrom 2008). Regulation of the actin cytoskeleton is critical for many biological and pathological events, including normal cell migration and metastasis. Indeed, Wrch-1 and Chp-1 have been shown to be important in focal adhesion turnover (Ory et al. 2007; Chuang et al. 2007; Aspenstrom et al. 2004), neural crest cell migration (Faure and Fort 2011; Notarnicola et al. 2008), and T-ALL cell migration (Bhavsar et al. 2013). Further studies should be aimed at examining the specific signaling pathways involved in CHW-1 axon guidance and neuronal migration.

To visualize the PDE neuron we utilized the *osm-6* promoter, which also allows visualization of the amphid neurons in the head and the phasmid neurons in the tail (Collet et al. 1998). When we visualized the PDE neuron, we observed that one of the GTPase mutants, CHW-1(A18G), resulted in altered amphid neuron morphology. The amphid neurons are born in the nose, and they migrate anteriorly to their final position where the bodies are located in the anterior region of the pharyngeal bulb and the axons that associate with the nerve ring. We noticed that the amphid neuron cell bodies in the CHW-1(A18G) animals were displaced anteriorly, similar to a loss of functions SAX-3 mutant (Zallen et al. 1999). In most Ras superfamily GTPases, the glycine at position 12 in Ras (position 18 in CHW-1) is highly conserved. Any other amino acid at this position is predicted to hinder the GTPase ability of the protein, rendering the protein at least partially activated. A valine at this position is the strongest activating mutation, resulting in a protein that has no GTPase activity (Wittinghofer et al. 1991) and is insensitive to GAPs, rendering the protein constitutively GTP-bound and active (Ahmadian et al. 1997). Normally, CHW-1 has an alanine at this position (position 18), which is predicted to be partially activating. Therefore, mutation of this alanine to a valine is predicated to be a stronger activation mutation, and mutation to a glycine would be predicted to decrease the activity of this protein. Indeed, when this alanine is mutated to a valine, it results in ectopic lamellipodia formation in the PDE neuron, similar to what is observed in CDC-42, CED-10, and MIG-2 (Shakir et al. 2008; Struckhoff and Lundquist 2003; Demarco and Lundquist). Furthermore, expression of the glycine mutant (CHW-1(A18G)) results in altered amphid neuron morphology, suggesting that CHW-1 activity is required for proper amphid neuron morphology and migration. CHW-1(A18G) may result in decreased function of the protein (by increased GTPase activity), but it also may act as a dominant negative. To further explore the role of CHW-1(A18G), we examined a null allele of *chw-1.* The null allele did not result in any defects in the amphid neurons, suggesting that CHW-1(A18G) is not acting simply as a protein with decreased function, and may be acting as a dominant negative. Further biochemical studies are needed to confirm this hypothesis.

This is the second study that examines GTPase mutants of CHW-1. In a paper by Kidd *et al.*, the authors showed that expression of wild-type CHW-1 resulted in errant distal tip cell (DTC) migration when compared to the GFP only negative control (Kidd et al. 2015). The authors also found that expression of CHW-1(A18G) abolished these defects, and expression of an activated CHW-1(A18V) mutant increased these defects, suggesting that native CHW-1 is partially activated. In our hands, ectopic expression of CHW-1 in the PDE neurons caused axon pathfinding defects, and neither GTPase mutant statistically altered the amount of defects, suggesting that in this system increased CHW-1 expression in this system is sufficient to induce axon guidance defects. We also found that activated CHW-1(A18V) induced ectopic lamellipodia in the PDE neurons, similar to the activated versions of other Rho proteins, supporting the findings of Kidd *et al.* In addition, we found the CHW-1(A18G) mutant may be a dominant negative mutation, complementing the previous findings.

With the CHW-1(A18G) mutant, we found that the amphid neurons were displaced anteriorly, similar to a *sax-3* loss of function phenotype. Therefore, we wanted to determine if *chw-1* interacted with *sax-3* in neuronal guidance. We found that *chw-1* interacts genetically with *sax-3* in amphid neurons, and in the PDE neurons. We found that loss of *chw-1* and *sax-3* synergistically increased the defects in amphid neurons and PDE axon pathfinding. These data suggest that these genes may be working in the same or redundant pathways. To further explore the relationship between *chw-1* and *sax-3*, we added an myristolyation site to *sax-3*, which results in an activated version of *sax-3* (*unc-25::myr-sax-3*). This modification has been previously used for other guidance receptors, such as UNC-40 (Gitai et al. 2003), to generate constitutively active receptors. Expression of MYR-SAX-3 in the VD/DD motor neurons resulted in axon pathfinding defects. These pathfinding defects were attenuated when we crossed in a null *chw-1* allele. These defects were not attenuated in the presence of a null *crp-1* allele. These data suggest that *chw-1* but not *crp-1* are working downstream of *sax-3* in axon pathfinding. These data were confirmed with RNAi, which showed that RNAi against *chw-1* but not *crp-1* decreased the axon guidance defects driven by activated SAX-3.

The *C. elegans* SAX-3/Robo is a guidance receptor, which acts in anterior-posterior, dorsal-ventral, and midline guidance decisions (Killeen and Sybingco 2008). Binding of the ligand Slit to the SAX-3/Robo receptor deactivates Cdc42 and activates another Rho GTPase, RhoA (Wong et al. 2001). In *Drosophila*, the adaptor protein Dock directly binds Robo and recruits p-21 activated kinase (Pak) and Sos, thus increasing Rac activity (Fan et al. 2003). Here, we show that there is a genetic interaction between *sax-3* and *chw-1*, and that *chw-1* is likely working downstream of *sax-3* in the VD/DD motor neurons. Activation of SAX-3 likely also results in signaling through CHW-1, in addition to other Rho GTPases, to mediate axon guidance. SAX-3/Robo signaling has also been implicated in carcinogenesis. Slit2 expression occurs in a large number of solid tumors, and Robo1 expression occurs in parallel in vascular endothelial cells. Concurrent expression of these proteins results in angiogenesis and increased tumor mass (Wang et al. 2003). Furthermore, mutations in SLT2, ROBO1, and ROBO2 are implicated in pancreatic ductal adenocarinoma (Biankin et al. 2012). Although the mammalian homologues Wrch-1 and Chp have been implicated in epithelial cell morphogenesis (Brady et al. 2009; Alan et al.), migration (Ory et al. 2007; Chuang et al. 2007; Faure and Fort 2011), and anchorage-independent growth (Berzat et al. 2005; Alan et al.; Chenette et al. 2006), the exact mechanisms by which these processes occur remains to be fully elucidated. One possible mechanism may be SAX-3/Robo signaling through CHW-1/Wrch-1. Further studies will be aimed at determining the precise mechanism by which these two molecules interact in both normal physiological processes and aberrant pathological processes.

In summary, we have shown that *chw-1* works redundantly with *crp-1* and *cdc-42* in axon guidance. Furthermore, proper levels and activity of CHW-1 are required for proper axon guidance in *C. elegans.* Importantly, we were able to show that CHW-1 is working downstream of the guidance receptor SAX-3/Robo. This novel pathway is important in axon guidance and may be important in development in addition to other pathogenic processes such as tumorigenesis, angiogenesis, and metastasis.

